# Multiple redundant mechanisms account for the majority of gene silencing downstream of DNA methylation

**DOI:** 10.64898/2026.02.08.704674

**Authors:** Shuya Wang, Zhongshou Wu, Zheng Li, Andrea Movilli, Li He, Yuxing Zhou, Evan K. Lin, Russell Chuang, Wai Wai Thiri, Sean Convery, Suhua Feng, Detlef Weigel, Steven E. Jacobsen

## Abstract

DNA methylation is a conserved epigenetic modification crucial for silencing genes and transposable elements (TEs). However, the mechanisms that cause silencing remain unclear, partly because methyl reader protein mutants in both plants and animals show minimal transcriptional changes. To explore the possibility of redundancy among these silencing mechanisms, we generated combinatorial mutants of H1.1, H1.2, ADCP1, MOM1, MBD2, MBD5, and MBD6 lacking key methyl readers and related silencing pathways. We observed massive derepression of genes and TEs at DNA-methylated loci, showing that these pathways account for 73% of silencing compared to DNA methylation-free mutants. We also observed that immune response genes were upregulated, causing an imbalance between growth and defense. Loss of downstream silencing pathways further disrupted 3D genome organization, leading to increased euchromatin–heterochromatin interactions. These findings highlight the cooperative action of multiple downstream mechanisms in DNA methylation-mediated silencing and genome organization.

## Introduction

DNA methylation, a conserved epigenetic modification, plays critical roles during development and diseases primarily via transcriptional silencing of genes and TEs [1-5]. DNA methylation can either block the access of transcriptional activators or recruit downstream readers with associated silencer complexes to repress transcription [6-14]. Despite efforts to understand the detailed silencing mechanisms downstream of DNA methylation, current studies have found little effect on transcription when these pathways are perturbed [6, 15]. For example, recent works from *Arabidopsis* have reported limited transcriptional changes at TEs in the mutant lacking the methyl readers *MBD2, MBD5*, and *MBD6* [14, 15]. MOM1 is another protein that binds to methylated regions of the genome and is required for transcriptional silencing, but its overall effects on gene expression are relatively small [16]. Mammalian studies have reached similar conclusions since an *MBD* quadruple mutant (*MBD1, MBD2, MBD4*, and *MeCP2*) exhibits mild changes in gene expression [6]. In contrast, dramatically more genes and TEs are strongly activated in DNA methylation-free mutants in *Arabidopsis* [17].

DNA methylation is known to have a significant impact on histone modifications. In *Arabidopsis*, dimethylated H3 lysine 9 (H3K9me2), a hallmark of silenced chromatin, is established and maintained via histone methyltransferases SUVH5, SUVH6, and KYP, which bind to methylated cytosine via their SRA domain [18]. As a result, the H3K9me2 modification pattern is primarily determined by the DNA methylation pattern. DNA methylation loss also leads to heterochromatin decompaction and affects the binding of linker histone H1 to heterochromatin [19, 20]. Perplexingly, H1 mutants, as well as mutations of a key reader of H3K9me2, ADCP1, also show very moderate effects on genome-wide transcription [21, 22]. The minimal transcriptional changes from disrupting each of the known silencing pathways related to DNA methylation suggest possible functional redundancy among them. Indeed, previous work has shown that knocking out linker histone H1 moderately increases transcriptional activation at genes and TEs in the *mbd5 mbd6* mutant [23], and a similar enhancement is observed in the *mbd2 mbd5 mbd6* mutant [14].

Here, we comprehensively examined the extent of redundancy among these pathways and quantified their overall contribution to the full range of DNA methylation-mediated silencing. We constructed an *h1*.*1 h1*.*2 adcp1 mom1 mbd2 mbd5 mbd6* (*hhammmm*) mutant, lacking the known methyl readers downstream of DNA methylation, together with mechanisms downstream of the epigenetic modifications that are lost in the DNA methylation-free mutant. The *hhammmm* mutants showed massive TE activation and gene activation relative to the single mutants, representing most of the upregulation seen in the DNA methylation-free mutants. TEs, including LTR/Gypsy, DNA/MuDR, and RC/Helitron, showed a prominent activation. Interestingly, such an upregulation was insufficient to trigger TE transposition as measured by both short-read and long-read sequencing. This indicates that the presence of additional redundant mechanisms, or possibly DNA methylation itself, regulates TE transposition. The *hhammmm* mutants also exhibited growth defects, likely caused by enhanced immune responses, illustrating how DNA methylation downstream mechanisms help balance the trade-offs between growth and immune defense. Micro-C showed increased interaction between euchromatin and heterochromatin in the *hhammmm* mutant, especially across the boundaries insulated by methyl readers. Finally, the *hhammmm* mutant transcriptional defects were still not as strong as those observed in mutants that were completely DNA methylation-free, suggesting that additional silencing pathways downstream of methylation likely exist. These results highlight that multiple redundant mechanisms contribute to gene and TE regulation by DNA methylation and in safeguarding the genome from excessive TE proliferation.

## Result

### DNA methylation remains intact despite the loss of downstream mechanisms

To assess the overall transcriptional effect of DNA methylation, we knocked out all five *Arabidopsis* methyltransferases, MET1, DRM1, DRM2, CMT2, and CMT3, generating the hereafter named *mddcc* mutant. We used CRISPR-Cas9 to knock out *MET1* in the *ddcc* (*drm1 drm2 cmt2 cmt3*) background and obtained the null mutant (fig. S1A). As a validation, we performed Whole-Genome Bisulfite Sequencing (WGBS) and found that the *mddcc* mutant exhibited a complete genome-wide erasure of CG, CHG, and CHH methylation (Fig. 1A-D and fig. S2A-B), which is consistent with the previously reported phenotype [17]. To evaluate redundancy of silencing pathways downstream of DNA methylation, we first knocked out the H3K9me2 reader *ADCP1* [22, 24] in the *h1*.*1 h1*.*2* T-DNA mutant background to create the *h1*.*1 h1*.*2 adcp1-3* triple mutant (Fig. S1B). Then we disrupted the additional silencing pathways downstream of DNA methylation, MOM1, MBD2, MBD5, and MBD6. We employed two genetic strategies (fig. S1C-H) and obtained two sets of alleles, referred to as *h1*.*1 h1*.*2 adcp1 mom1 mbd2 mbd5 mbd6-1 (hhammmm-1)* and *hhammmm-2* mutants.

**Fig. 1.**
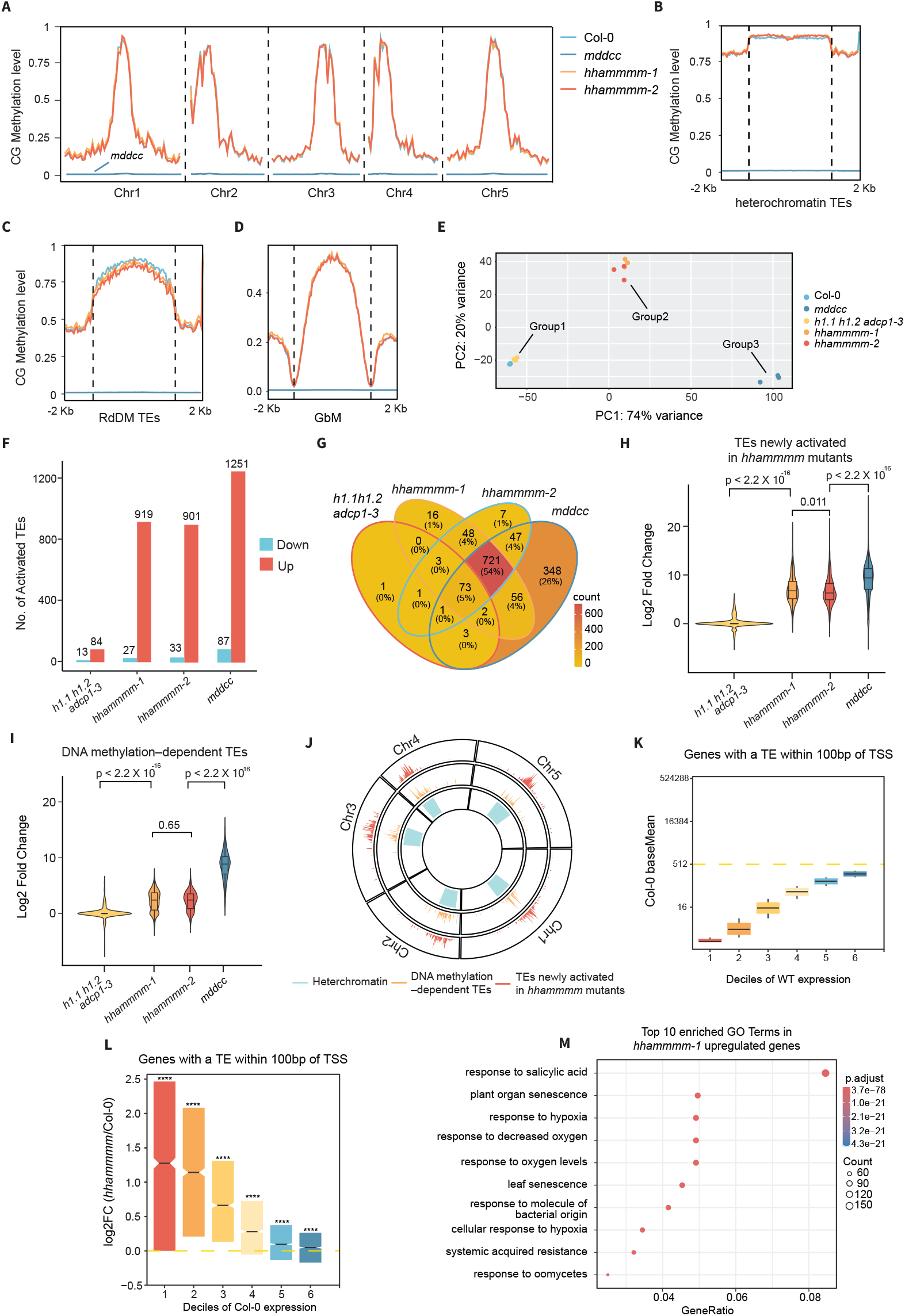
Mechanisms downstream of DNA methylation play a key role in silencing genes and TEs. **A**. Line plot showing the genome-wide CG methylation level in Col-0, *mddcc, hhammmm-1*, and *hhammmm-2* mutants. Metaplots demonstrating CG methylation level at **B**. heterochromatic TEs, **C**. RdDM TEs, and **D**. genes with gene-body methylation (GbM). **E**. PCA analysis based on TE transcriptomics of Col-0, *mddcc, hhammmm-1*, and *hhammmm-2* mutants. Three groups are defined by sample proximity. **F**. Bar plot displaying the number of differentially expressed TEs in *h1*.*1 h1*.*2 adcp1-3, hhammmm-1, hhammmm-2*, and *mddcc* mutants. **G**. Venn diagram illustrating the overlaps among the significantly activated TEs in *h1*.*1 h1*.*2 adcp1-3, hhammmm-1, hhammmm-2*, and *mddcc* mutants. Violin and box plots demonstrating the log_2_ fold change (FC) of TE expression at **H**. TEs newly activated in the *hhammmm* mutants, or **I**. TEs only activated in the *mddcc* mutant. **J**. Circle plot showing the genome-wide distribution of activated TEs. Heterochromatin is defined by enrichment of H3K9me2. **K**. Box plot showing the average basal expression of six gene deciles. Only genes having an activated TE within 100 bp of TSS are considered. **L**. Box plot displaying the log_2_FC of gene expression in the *hhammmm* mutants compared to Col-0 across six gene deciles. **M**. Top 10 enriched GO terms of *hhammmm-1* upregulated genes.

To test the effects of these downstream pathways on DNA methylation, we performed WGBS for the *hhammmm* mutants, with wild-type (Col-0) and the *mddcc* mutant as controls. Wild-type and the *hhammmm* mutants showed similar CG methylation levels genome-wide, while the *mddcc* mutant was completely free of DNA methylation (Fig. 1A). We also examined the CG methylation at long heterochromatic TEs (Fig. 1B), RNA-directed DNA methylation (RdDM) targeted TEs (Fig. 1C), and genes with gene-body methylation (GbM) (Fig. 1D). The *hhammmm* mutants exhibited similar CG methylation levels to those of the wild-type at all genomic features examined (Fig. 1A-D). As for non-CG methylation, we found a slight increase in CHG methylation at heterochromatin regions (fig. S2A-B). Our observation is consistent with the previously reported observation that in the *h1*.*1 h1*.*2* mutant, CMT3 gains more access to heterochromatic nucleosomes [25, 26]. Meanwhile, CHH methylation was slightly reduced in heterochromatin (fig. S2B), which likely resulted from ADCP1 loss as observed previously [22]. Although we observed a mild increase in CHG and a slight decrease in CHH methylation, the *hhammmm* mutants overall preserved the majority of DNA methylation across all sequence contexts, thus allowing for examination of the silencing capacity of DNA methylation and its downstream mechanisms.

### Downstream mechanisms account for most silencing activity caused by DNA methylation

We then carried out RNA sequencing (RNA-seq) to evaluate transcriptional activation in wild-type, *h1*.*1 h1*.*2 adcp1-3, hhammmm-1, hhammmm-2*, and *mddcc* mutants. We focused our analysis primarily on TEs to quantify the direct influences on transcriptional silencing, since many changes in protein-coding genes may come from indirect effects [27]. From the Principal Component Analysis (PCA) based on TE expression, the four genotypes tested clustered into three distinct groups (Fig. 1E). Col-0 and *h1*.*1 h1*.*2 adcp1-3* samples clustered together and were defined as group1, suggesting that *h1*.*1 h1*.*2 adcp1-3* leads to a limited level of TE activation. The two independent *hhammmm* mutants formed group2 and clustered away from group1. The *mddcc* mutant clustered as the third group and was distinct from the *h1*.*1 h1*.*2 adcp1-3* and the *hhammmm* mutants (Fig. 1E). To further assess the level of TE activation in these mutants, we examined the number of significantly upregulated TEs. There were 84 TEs de-repressed in *h1*.*1 h1*.*2 adcp1-3* mutant (Fig. 1F), representing a minimal level of transcriptional activation of TEs. In the *hhammmm* mutants, we observed that more than 900 TEs were upregulated, more than a 10-fold increase compared to the *h1*.*1 h1*.*2 adcp1-3* triple mutant (Fig. 1F). Erasure of DNA methylation in the *mddcc* mutant caused transcriptional activation of 1251 TEs, around 300 more than in the *hhammmm* mutants (Fig. 1F-G and fig. S3A), suggesting that DNA methylation itself and/or additional downstream silencing pathways contribute to the full strength of TE silencing. To more quantitatively assess the contributions of DNA methylation and its downstream mechanisms to transcriptional silencing, we compared the expression levels of derepressed TEs, mainly focusing on two groups of upregulated TEs: (i) TEs that are newly activated in the *hhammmm* mutants compared to the *h1*.*1 h1*.*2 adcp1-3* mutant, and (ii) TEs that are only significantly upregulated in the *mddcc* methylation-free mutant (Fig. 1G). In the first group, loss of *MOM1* and *MBDs* caused a 32-fold increase in the level of TE transcription, and loss of DNA methylation further enhanced this effect by another 16-fold (Fig. 1H and fig. S3B). These results show that there is substantial redundancy in the degree of transcriptional repression controlled by the MBDs, MOM1, ADCP1, and H1 linker histone. They also highlight that the methylation-free mutant exhibits greater TE upregulation compared to the *hhammmm* mutants, suggesting that additional downstream silencing pathways must be important. Upon the examination of group2 TEs, which were only significantly upregulated in the *mddcc* mutant (Fig. 1G and fig. S3A), we discovered that they were already mildly activated in the *hhammmm* mutants (Fig. 1I and fig. S3B). These data suggest that almost all TEs silenced by DNA methylation are derepressed to some extent in the *hhammmm* mutant, with different levels of activation among TEs. In addition, the upregulated TEs in the *mddcc* and the *hhammmm* mutants were concentrated in the pericentromeric heterochromatic regions where DNA methylation density is the highest (Fig. 1J). To further validate that the observed TE activation is not a result of local loss of DNA methylation, we examined the CG methylation level at the group1 TEs, which are newly activated in the *hhammmm* mutants. The *hhammmm* mutants exhibited no major changes in DNA methylation at these TEs, with only a small subset displaying mild hypomethylation (fig. S3C), which is likely caused by transcription-coupled DNA demethylation [28]. Taken together, our results suggest that several known silencing pathways downstream of DNA methylation function cooperatively/redundantly to exert the full extent of TE silencing, and a substantial fraction of TEs are only derepressed when multiple layers of these silencing mechanisms are simultaneously disrupted. In addition to silencing TE transcription, DNA methylation downstream pathways are also required for regulating the expression of protein-coding genes [17]. For example, the upregulation of TEs can affect neighboring gene expression [29-31]. Indeed, in *hhammmm* mutants, genes with their promoters either containing or proximal (within 100bp) to the activated TEs exhibited significant upregulation, especially for genes with relatively lower expression in wild-type plants (Fig. 1K-L). We also performed Gene Ontology (GO) analysis for significantly upregulated genes in the *hhammmm* mutants and found that the upregulated genes were enriched in the GO terms of salicylic acid-mediated defense (Fig. 1M and fig. S4A). Consistent with this GO term enrichment, the *hhammmm* mutants have phenotypes resembling those of classical autoimmune mutants, such as reduced size and curly leaves (fig. S4B-C)[32, 33]. Consistent with previous work [17], immune response-related GO terms were also significantly enriched in the *mddcc* mutant (fig. S4D), and extreme developmental and reproductive defects were observed (fig. S4B-C) [34].

*Arabidopsis* TEs are classified into different families based on their transposition mechanisms and internal structures. We analyzed the TE family profile of the TEs upregulated in the *hhammmm* mutants, as well as the TEs that are only significantly upregulated in the *mddcc* mutant. Both groups of TEs exhibited the most enrichment in the LTR/Gypsy family, followed by the DNA/En-Spm and the DNA/MuDR families (Fig. 2A). LTR/Gypsy TEs also displayed a slightly higher degree of activation compared to other TE families in the *hhammmm* mutants while DNA/En-Spm TEs showed the most prominent activation in the *mddcc* mutant, suggesting that DNA methylation and related downstream pathways have different strengths in silencing specific TE families (Fig. 2B). A small fraction of LTR/Gypsy TEs can transpose in certain genetic backgrounds and/or environmental conditions. For example, heat stress induces the transposition of ONSEN and DNA hypomethylation triggers the transposition of EVADE [35, 36]. Since DNA methylation and downstream pathways repress the transcription of LTR/Gypsy, DNA/En-Spm, and DNA/MuDR, we tested whether increased TE expression in the *hhammmm* mutants was sufficient to trigger TE transpositions. To determine this, we used long-read sequencing to assess the extent of somatic transposition in Col-0, *h1*.*1 h1*.*2 adcp1-3*, and the *hhammmm* mutants. We observed a similarly low number of somatic transpositions across all genotypes (fig. S5A-C). The extent of somatic mobilization we found in our mutants was substantially lower than that reported for *met1* mutants of the Tsu-0 accession background, using the same long-read approach (fig. S5A-C) [37]. Complementary results from whole-genome sequencing (WGS) using short reads detected TE transposition events only in *mddcc* mutants but not in *h1*.*1 h1*.*2 adcp1-3* or *hhammmm* mutants (fig. S5D-E). Since *ddm1, met1 cmt3*, as well as *met1*, also cause a much higher level of TE transposition [38], DNA methylation itself appears to play a prominent role in preventing TE transposition, likely via physically obstructing transposase-DNA interactions at the binding sites [39], or there may be additional downstream silencing mechanisms that are critical.

**Fig. 2.**
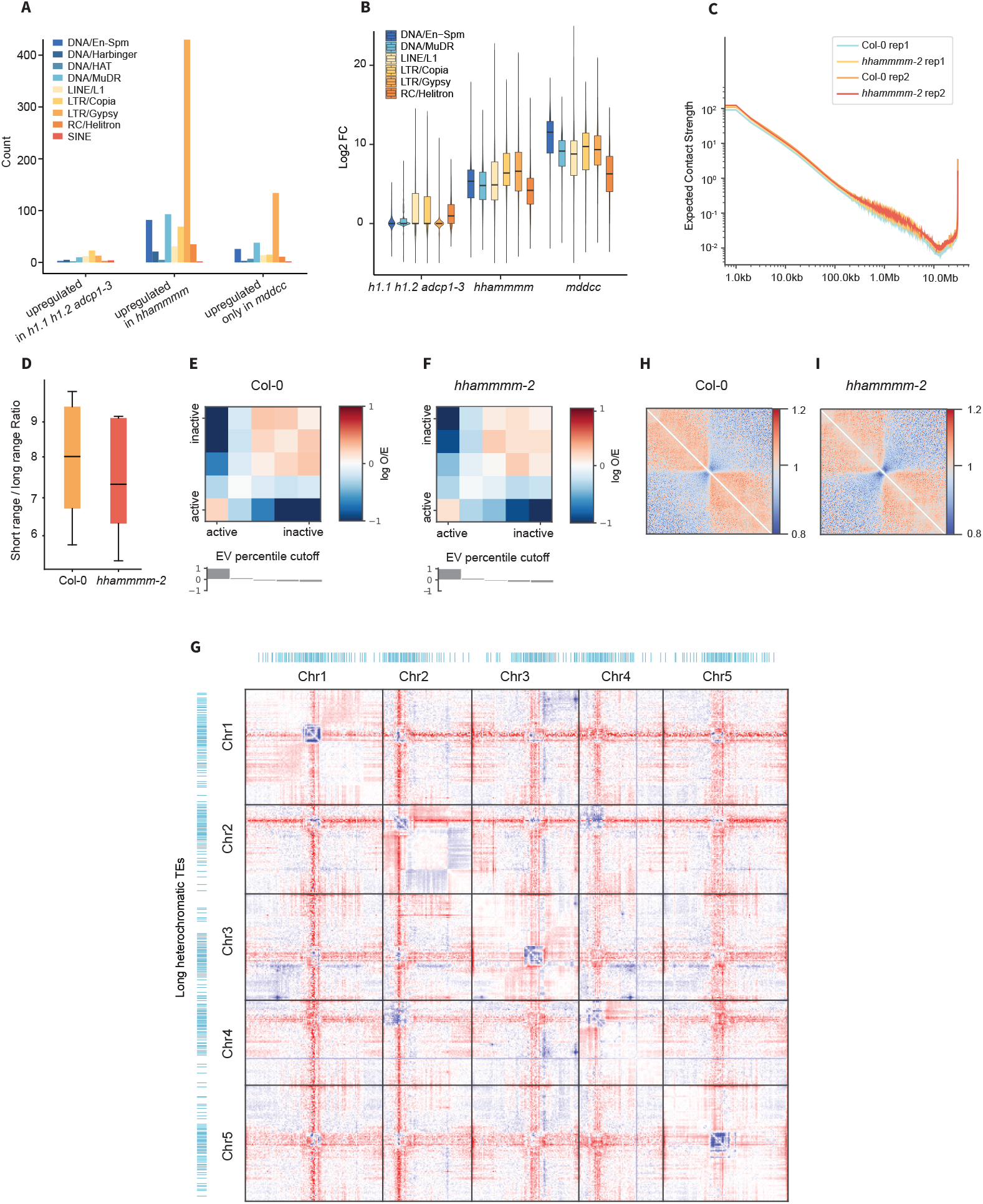
Downstream mechanisms ensure proper 3D genome folding. **A**. Bar plot demonstrating the number of activated TEs under 9 TE families. Three TE groups are listed: TEs upregulated in *h1*.*1 h1*.*2 adcp1-3, hhammmm*, or *mddcc* mutants. **B**. Box plot showing the log_2_FC of TEs from three TE groups. For each group, gene expression changes of the top six upregulated TE families were plotted. **C**. Distance-dependent contact strength plot of Col-0 and *hhammmm-2* mutant. **D**. Box plot showing the ratio of short-range versus long-range interactions between Col-0 and *hhammmm-2* mutant. **E**. and **F**. Matrices illustrating the observed/expected contact frequency together with eigen vector plots between A and B compartments of Col-0 (E), and *hhammmm-2* mutant (F). **G**. Micro-C map showing the differences of the interactions between *hhammmm-2* and Col-0 at 500 Kb resolution. **H**. and **I**. Aggregate analysis of chromatin interactions at euchromatic islands within the pericentromeric heterochromatic regions for Col-0 (H) and *hhammmm-2* (I).

### Downstream mechanisms of DNA methylation influence 3D genome organization

Loss of DNA methylation has been shown to disrupt normal 3D genome organization [40, 41]. Considering the similarity of the overall TE activation landscapes of the *hhammmm* and *mddcc* mutants, we hypothesized that the 3D genome architecture would also be altered in the *hhammmm* mutants. To test this, we applied Micro-C to compare chromatin folding between wild-type and the *hhammmm* mutants at nucleosome-level resolution. As a validation of the methodology, the Micro-C technique successfully reproduced previously reported chromatin features, including compartments, Interactive Heterochromatic Islands (IHIs), and chromatin loops (Fig. 2C–I, Table S1)[40, 42]. In the *hhammmm-1* mutant, we detected a translocation between chromosomes 3 and 5, probably resulting from T-DNA inserting into the *MBD6* gene on chromosome 5 and a coincident double-strand break in the arm of chromosome 3 (fig. S6A). This is a common phenomenon in T-DNA mutants [43, 44]. To avoid potential misinterpretation of Micro-C data caused by this translocation, we focused subsequent analyses on the *hhammmm-2* mutant. We found that the *hhammmm-2* mutant exhibits more frequent long-range (>1 Mb) chromatin contacts compared to the wild-type, indicating reduced insulation among chromatin domains (Fig. 2C–D). Furthermore, A and B compartments in the *hhammmm-2* mutant showed increased inter-compartmental interactions, both within and between chromosomes (Fig. 2E–G, fig. S6B-D). These changes were more pronounced than those previously observed in the *h1*.*1 h1*.*2* mutants, especially for inter-chromosomal interactions [45]. Because the active A and inactive B compartments show more frequent interactions in the *hhammmm-2* mutant, we performed a more detailed analysis focusing on the euchromatic islands within the pericentromeric heterochromatic regions. In the wild-type, local heterochromatic domains on each side of euchromatic patches displayed a low level of interaction, suggesting insulation. In the *hhammmm-2* mutant, euchromatic islands showed more interaction with distal heterochromatin, causing erosion of local heterochromatin boundaries (Fig. 2H-I). Overall, our data demonstrate that mechanisms downstream of DNA methylation help define the limits of both chromosome-scale A/B compartments and local gene-level domains, thereby shaping chromatin compartmentalization.

## Discussion

While the fundamental role of DNA methylation in transcriptional silencing has been known for decades, the critical downstream mechanisms of silencing have remained understudied. Mutations of downstream silencing components, especially those that preserve DNA methylation, have thus far failed to recapitulate the transcriptional changes caused by the loss of DNA methylation. This conundrum has led to extensive efforts to identify the various silencing pathways downstream of DNA methylation, as well as controversies about whether prominent DNA methylation-dependent silencing pathways even exist, and whether DNA methylation mainly acts to directly block the binding of transcription factors to methylated DNA [6, 15]. Our findings unequivocally demonstrate that downstream silencing mechanisms are indispensable for DNA methylation-mediated transcriptional silencing, and show that virtually all TEs silenced by DNA methylation are also silenced to some extent by downstream silencing. . The pervasive TE upregulation in the *hhammmm* mutants compared to previously studied mutations in single downstream silencing pathways suggests that substantial redundant action among these pathways is pivotal for the robust silencing exerted by DNA methylation. The redundant action of these different pathways not only offers extra layers of protection against excessive TE activation but also likely provides different fine-tuning handles for precise transcriptional control. An interesting question is why there is so much redundancy in silencing mechanisms downstream of methylation. One possible explanation is that intense evolutionary pressure from TE expansions of different TE types may have facilitated the evolution of many different countermeasures to silence TEs.

Although the *hhammmm* mutants accounted for the majority of TE activation observed in the *mddcc* mutant, residual differences in both the magnitude and quantity of activated TEs indicate that additional silencing pathways and/or DNA methylation itself act to repress transcription. For example, other MBD proteins [46], MORC proteins [47], and ADCP2 [48] may play important roles in silencing TEs and act redundantly with the pathways described in this study. Furthermore, DNA methylation can directly prevent transcription factor binding, and zinc-finger-type transcription factors are known to sometimes recognize methylated cytosine as thymine [49]. These mechanisms can alter transcription factor activity, thereby affecting TE silencing. The detailed contribution of transcriptional regulation by DNA methylation itself versus that of downstream pathways requires evaluation in future studies.

Beyond TE silencing, we found that pathways acting downstream of DNA methylation help define 3D chromatin compartment boundaries by serving as insulators in both heterochromatic and euchromatic regions. Moreover, our data suggest that these downstream DNA methylation pathways also regulate protein-coding gene expression, especially those genes with TEs near their start sites.

Collectively, our results highlight the crucial role of downstream repressive mechanisms in maintaining transcriptional silencing, preserving genome architecture, and modulating immune defense to protect plant development.

## Acknowledgments

We thank Yan He and Christian Fonkalsrud for technical support, Colette L. Picard for scientific discussion, and Mahnaz Akhavan, Suhua Feng, and the UCLA BSCRC BioSequencing Core for sequencing assistance.

## Funding

National Institutes of Health grant R35 GM130272 (SEJ)

Life Sciences Research Foundation postdoctoral fellowship (ZW)

Howard Hughes Medical Institute (SEJ)

## Author contributions

Conceptualization: SW, ZL, ZW, SEJ

Methodology: SW, ZW, ZL, AM, LH, YZ, EKL, RC, WWT, SC, SF

Investigation: SW, ZW, ZL, SEJ, AM, LH, DW

Funding acquisition: ZW, SEJ

Supervision: SW, ZW, ZL, SEJ

Writing – original draft: SW, ZW, ZL, SEJ

Writing – review & editing: SW, ZW, ZL, SEJ, AM, LH, DW

## Competing interests

S.E.J is the scientific co-founder of Inari Agriculture and a consultant for Inari, Invaio Sciences, Terrana Biosciences, Sail Biomedicines, and Zymo Research.

## Data and materials availability

All high-throughput sequencing data generated in this study will be publicly available.

## Supplementary Materials

Materials and Methods

Figs. S1 to S6

Tables S1

References (*14, 32, 44-52*)

