## Supplementary data for "Multiple redundant mechanisms account for the majority of gene silencing downstream of DNA methylation"

**The PDF file includes:**

Materials and Methods
Figs. S1 to S6
Tables S1

### Materials and Methods

Plant materials and growth conditions

The plants used in this paper were *Arabidopsis thaliana* Col-0 ecotype and were grown under long-day conditions (16 h light and 8 h dark). The T-DNA insertion lines used in this study include *h1.1–1* (*SALK_128430*), *h1.2–1* (*GABI_406H11*), and *mbd6* (*SALK_043927*). CRISPR constructs were generated based on the pHEE401E or pBEE401 vectors, based on a previously described CRISPR–Cas9 gene editing system [44]. To create the *mddcc* mutant, a CRISPR construct targeting MET1 was transformed into the *ddcc* mutant. The guide sequence used was ACCTAGGACGAGGAGACCA. We named the *met1* allele generated in the *ddcc* background *met1-10*, which carries a 2bp deletion near the guide target that resulted in a premature stop codon in its first exon. The *h1.1 h1.2 adcp1-3* triple mutant was generated by transforming the ADCP1 CRISPR construct into the *h1.1 h1.2* double mutant. The guide sequences used were CACTTGCAACGCCATGTTG and CAGTCTCAGAAGGTGTAAA, resulting in a 1953bp deletion, nearly removing the entire gene. To generate additional intermediate mutants, we started by knocking out *MOM1* in the *h1.1 h1.2* double mutant via CRISPR. The guides used for this transformation were ACTTTCTAGTAGCAGAAGC and GACGAGGTTGGAGTAGCTG, resulting in an 8212bp deletion in the *MOM1* gene. This newly generated *mom1* allele was designated as *mom1-5* (Fig. S1). Next, we knocked out both *MBD2* and *ADCP1* via CRISPR in the *h1.1 h1.2 mom1-5* mutant background using the same guides mentioned above. This approach produced the intermediate *h1.1 h1.2 mom1-5 mbd2-2 adcp1-4* quintuple mutant (Fig. S1). To generate the *h1.1 h1.2 mom1 mbd2 mbd5 mbd6 adcp1* septuple mutants, referred to as *hhammmm* mutants, we employed two strategies. *hhammmm-1* was obtained by crossing the *h1.1 h1.2 mom1-5* mutant with the reported *mbd2-1 mbd5-1 mbd6-1* triple mutant [14]. This triple mutant also carried the aforementioned ADCP1 CRISPR construct, which introduced the *adcp1-3* mutation in the F1 generation. Alternatively, *hhammmm-2* was generated by transformation of a CRISPR construct targeting both *MBD5* and *MBD6* genes in *h1.1 h1.2 mom1-5 mbd2-2 adcp1-4* quintuple mutant. MBD5 guides included TCTGATTCAGTCAAAAAGAA and CTGATCGTTGGTTGGAATGC, which introduced a 1111bp deletion, while MBD6 guide GTGCTGTACTACTTGGAACA caused a 7bp deletion that resulted in an early stop codon. The resulting *MBD5* and *MBD6* alleles were named *mbd5-2* and *mbd6-2*, respectively.

Whole genome bisulfite sequencing (WGBS)

DNA from 14-day-old seedlings was extracted using the Qiagen DNeasy Plant Mini Kit (Qiagen 69106). RNA was removed with PureLink RNase A (Invitrogen). A total of 100 ng of DNA was sheared to 200 bp (Duty Cycle = 10%, Intensity = 5, Cycles per Burst = 200, Treatment time = 120 seconds) with a Covaris S2 (Covaris). Libraries were prepared using the Epitect Bisulfite Conversion kit (QIAGEN) and the Ovation Ultralow Methyl-seq kit (NuGEN) according to the manufacturer's instructions. The libraries were sequenced on Illumina NovaSeq X Plus instruments.

RNA-seq

Six biological replicates were used for each genotype for RNA-seq. 14-day-old seedlings were harvested and frozen in liquid nitrogen. The samples were ground into powder, and RNA was extracted using the Direct-zol RNA MiniPrep kit (Zymo Research). 250 ng of total RNA were used for RNA-seq library preparation with the TruSeq Stranded mRNA kit (Illumina). The final library was sequenced on Illumina NovaSeq X Plus instruments.

Whole genome sequencing (WGS)

DNA was extracted and sheared using the same methods as in WGBS. Libraries were prepared with the Ovation Ultra Low System V2 kit (NuGEN) and sequenced on Illumina NovaSeq X Plus instruments.

HiFi library preparation

High-molecular-weight (HMW) DNA from pooled plant material of *h1.1 h1.2 adcp1-3*, *hhammmm-1*, *hhammmm-2*, and Col-0 wild-type was extracted following the published protocol (https://doi.org/10.2144/000114460). HMW DNA was then sheared using Megaruptor 2 (Diagenode). Sheared DNA was used as input for preparing HiFi SMRTbell libraries with the SMRTbell Express Template Prep Kit 3.0 (PacBio). Libraries were size-selected on a BluePippin (SageScience) using a 10 kb cutoff in a 0.75% DF Marker S1 High Pass 6-10 kb v3 gel cassette (Biozym). Final libraries were loaded onto Revio SMRT Cell 25M chips and sequenced on the PacBio Revio system using the Binding Kit 3.2 (PacBio).

Micro-C

1.5 g of 14-day-old seedlings were harvested and flash frozen. Samples were then ground into powder and dissolved in Nuclear Isolation Buffer (300 mM Sucrose, 20 mM Tris pH 8.0, 5 mM $MgCl_2$, 5 mM KCl, 0.2% Triton X-100, 35% Glycerol) for sonication. Sonicated samples, which supposedly contain nuclei, were subject to 30 min 300 mM DSG crosslinking followed by 10 min 1% Formaldehyde treatment. The crosslinking process was quenched by 128 mM Glycine. Samples were passed through two layers of Miracloth, and nuclei were purified through two rounds of buffer wash (Extraction Buffer II (0.25 M sucrose, 10 mM Tris-HCl pH 8.0, 10 mM MgCl2, 1% Triton X-100) and Extraction Buffer III (1.7 M sucrose, 10 mM Tris-HCl pH 8.0, 2 mM $MgCl_2$, 0.15% Triton X-100)). Nuclei were then rinsed with digestion buffer (0.32 M sucrose, 50 mM Tris (8.0), 4 mM $MgCl_2$, 1 mM $CaCl_2$), and then MNase (final concentration 0.2 U/ul) was added to digest chromatin into mono-nucleosomes. The reaction typically takes 30 min at 37 ℃, but should be optimized before each experiment. The reaction was stopped by incubating for 10 minutes at 65 ℃ with 10 mM EDTA and EGTA. Next, samples were end-repaired, end-chewed, and end-labeled with Biotin-dATP/dCTP and dGTP/dTTP. Afterwards, proximity ligation was done and bio-dNTP was removed at the unligated ends. Finally, the samples were reverse-crosslinked, and the DNA was purified. Library preparation was performed using the Kapa Hyper Prep kit, coupled with a streptavidin bead pull-down to enrich the ligated population before PCR amplification.

WGBS analysis

WGBS data were filtered, and Illumina adaptors were trimmed using Trim Galore (v 0.6.7, Babraham Institute). Reads containing three or more consecutive methylated CHH sites were classified as non-converted and excluded from analysis. Bismark (v 0.19.1, Babraham Institute)[45] was employed to align reads to the *Arabidopsis* reference genome (TAIR10). ViewBS (v 0.1.11)[46] generated the visual plots. DNA methylation levels were initially calculated and visualized using deepTools (v3.0.2)[47] with the computeMatrix and plotHeatmap functions.

RNA-seq analysis

The bowtie2 version 2.3.4.3 [48] was used to align the raw reads of RNA-seq data to the TAIR10 transcriptome, and rsem-calculate-expression of RSEM was used to calculate the expression levels with default settings [49]. The Samtools version 1.9 [50] was used to generate tracks, and the bamCoverage of deeptools version 3.1.3 was used to normalize the data with RPKM. DESeq2 (v.1.42.0) was used to perform the differential analysis with the cut-offs Padj $< 0.05$ and $|\log2FC| \geq 1$. Data presented in box plots have been normalized to the wild-type. We used ggplot2 (v.3.4.4) to generate all the related plots. We took the union of the activated TEs from mutants to generate the box plots.

WGS analysis

WGS data obtained from Col-0, *mddcc*, and *hhammmm* mutants were analyzed to detect non-reference TE insertions using ngs_te_mapper2 [51] (https://github.com/bergmanlab/ngs_te_mapper2) and TEFLoN [52] (https://github.com/jradrion/TEFLoN) with the TAIR10 reference genome and default parameters. To minimize false positive events, TE insertions supported by only a single read end (either 3′ or 5′) or detected by only one pipeline (ngs_te_mapper2 or TEFLoN) were excluded from further analysis. For each detected TE insertion, WGS reads were aligned to a custom pseudo-genome—comprising both the reference genome sequence and the inserted TE sequence—using BWA mem (v0.7.17). The resulting BAM files were then loaded into IGV for manual inspection of the insertion events.

Somatic transposition calling

Somatic TE insertions were identified and visually confirmed using the IGV genome browser, following the approach described in Movilli et al [32]. Reads from mutants and wild-type samples were aligned to the TAIR12 (Col-CC) reference genome, and the concordance with the TAIR12 TE annotation (https://github.com/oushujun/TAIR12-TE) was leveraged to call low-coverage insertions. We discarded all the events involving satellites and rDNA rearrangements due to their associated intrinsically low confidence. Insertions supported by reads that did not span the full length of the transposed TE, i.e., single-ended support, were classified as “low confidence”. In contrast, insertions with reads spanning the entire TE (both-ended support), were considered “high confidence”. For the latter case, we confirmed the presence of the associated target site duplication (TSD) as described in Movilli et al. A TE associated with one “high-confidence” event was annotated as three adjacent elements, but it appeared to mobilize as one. We therefore renamed it ‘ATMU13 composite’.

Micro-C analysis

Micro-C analysis was performed using the Dovetail Genomics pipeline (https://micro-c.readthedocs.io). First, read pairs were mapped to the TAIR10 genome using BWA-MEM (https://github.com/lh3/bwa). Ligation junctions were then identified, and PCR duplicates along with reads of low mapping quality were removed using pairtools (https://github.com/open2c/pairtools). The contact matrices were visualized with Juicer box. Contact information stored in mcool files was extracted with Cooler (https://github.com/open2c/cooler) and subsequently normalized using Cooler balance. Distancedependent decay of contact density plots was generated through FANC, and insulation score analysis was also conducted using FANC tools.

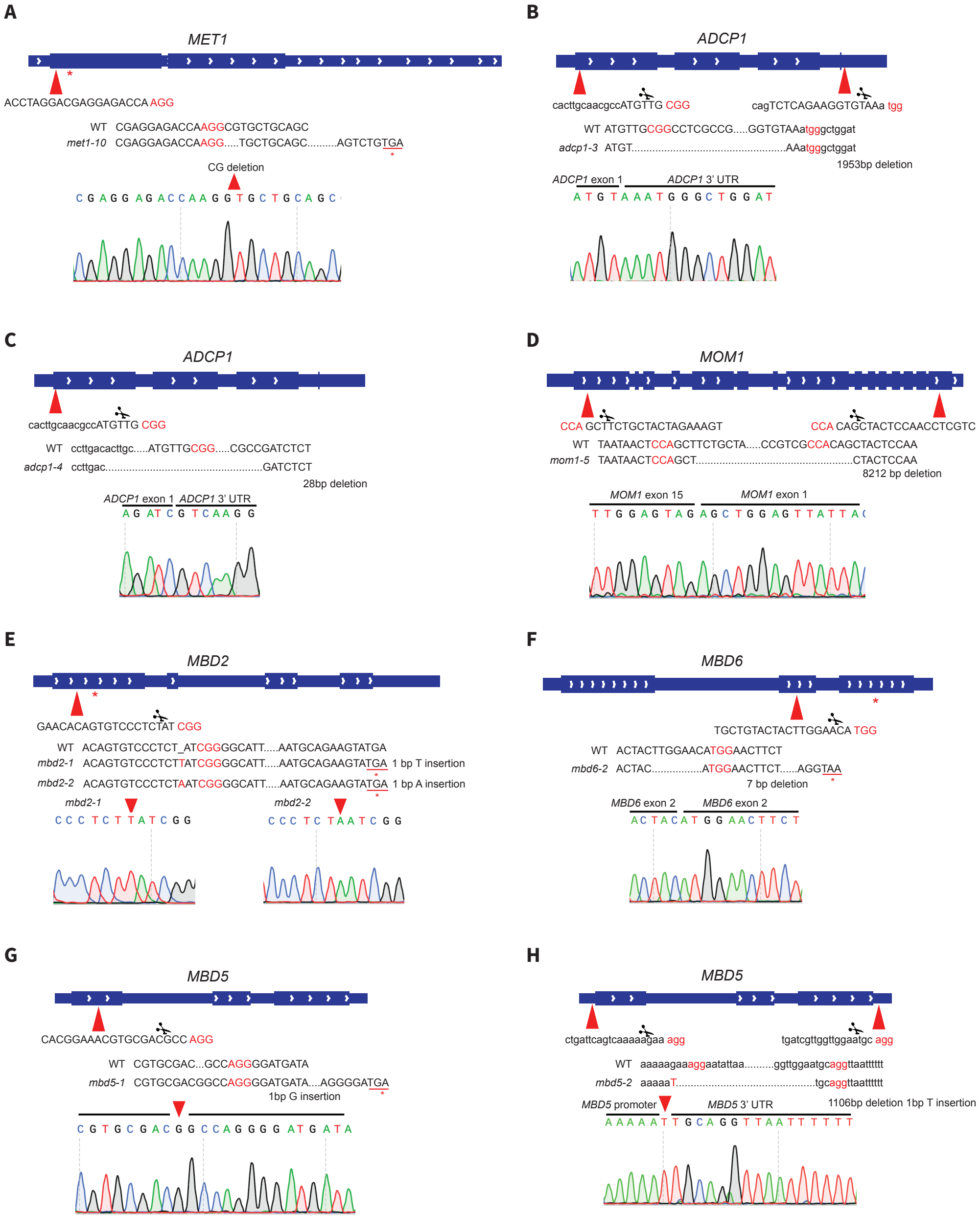


**Fig. S1. Confirmation of CRISPR-Cas9-generated mutations.**

Annotations and Sanger sequencing results showing the CRISPR-Cas9-mediated mutations at MET1 (**A**), ACDP1 allele 3 (**B**), ADCP1 allele 4 (**C**), MOM1 (**D**), MBD2 (**E**), MBD6 (**F**), MBD5 allele 1 (**G**), and MBD5 allele 2 (**H**).

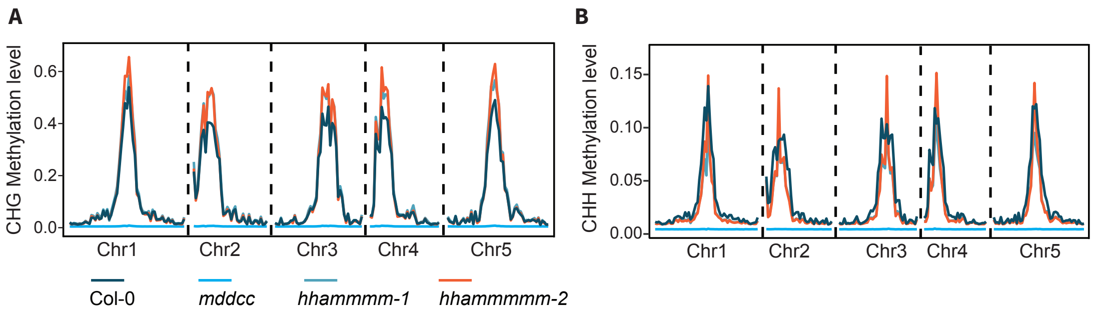


**Fig. S2. Non-CG methylation underwent minor changes in the *hhammmm* mutants.**
Line plots showing the genome-wide **A.** CHG methylation, and **B.** CHH methylation level of Col-0, *mddcc*, *hhammmm-1*, and *hhammmm-2* mutants.

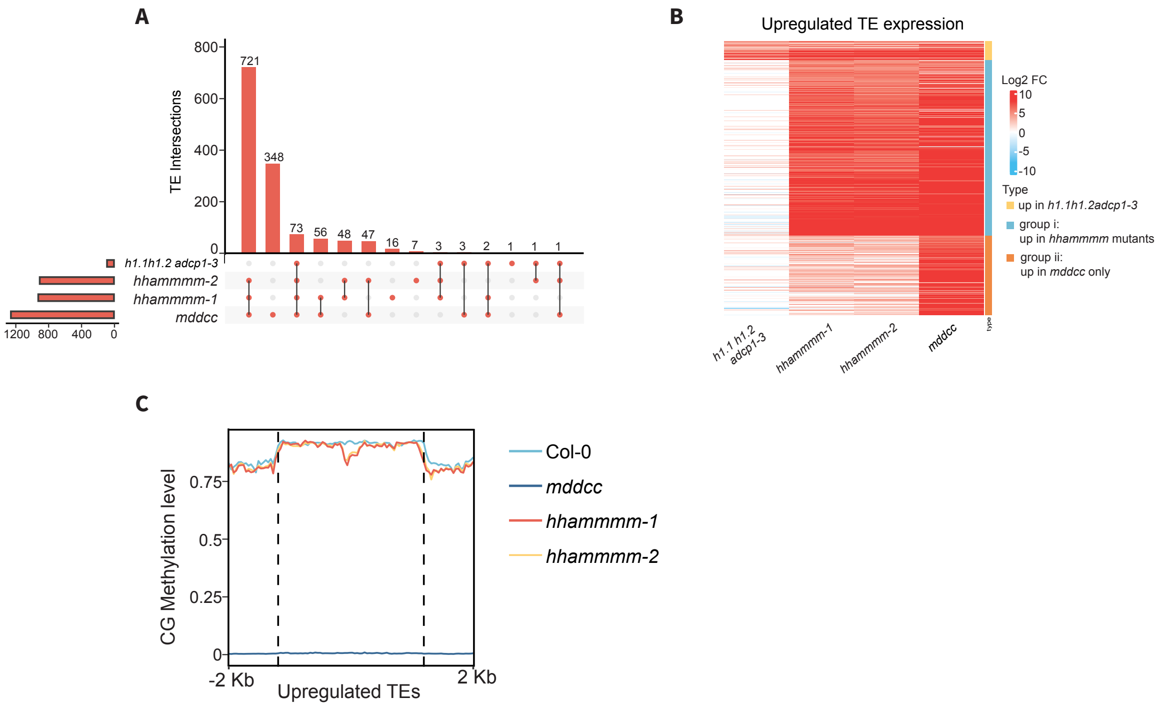


**Fig. S3. Loss of downstream mechanisms leads to a significant upregulation of TE without affecting the DNA methylation level.**

**A.** Upset plot showing the distribution of upregulated TEs from *h1.1 h1.2 adcp1-3*, *hhammmm*, and *mddcc* mutants. **B.** Heatmap showing the $\log_2$FC of upregulated TEs from *h1.1 h1.2 adcp1-3*, *hhammmm*, and *mddcc* mutants. **C.** Metaplot demonstrating the CG methylation level at upregulated TEs.

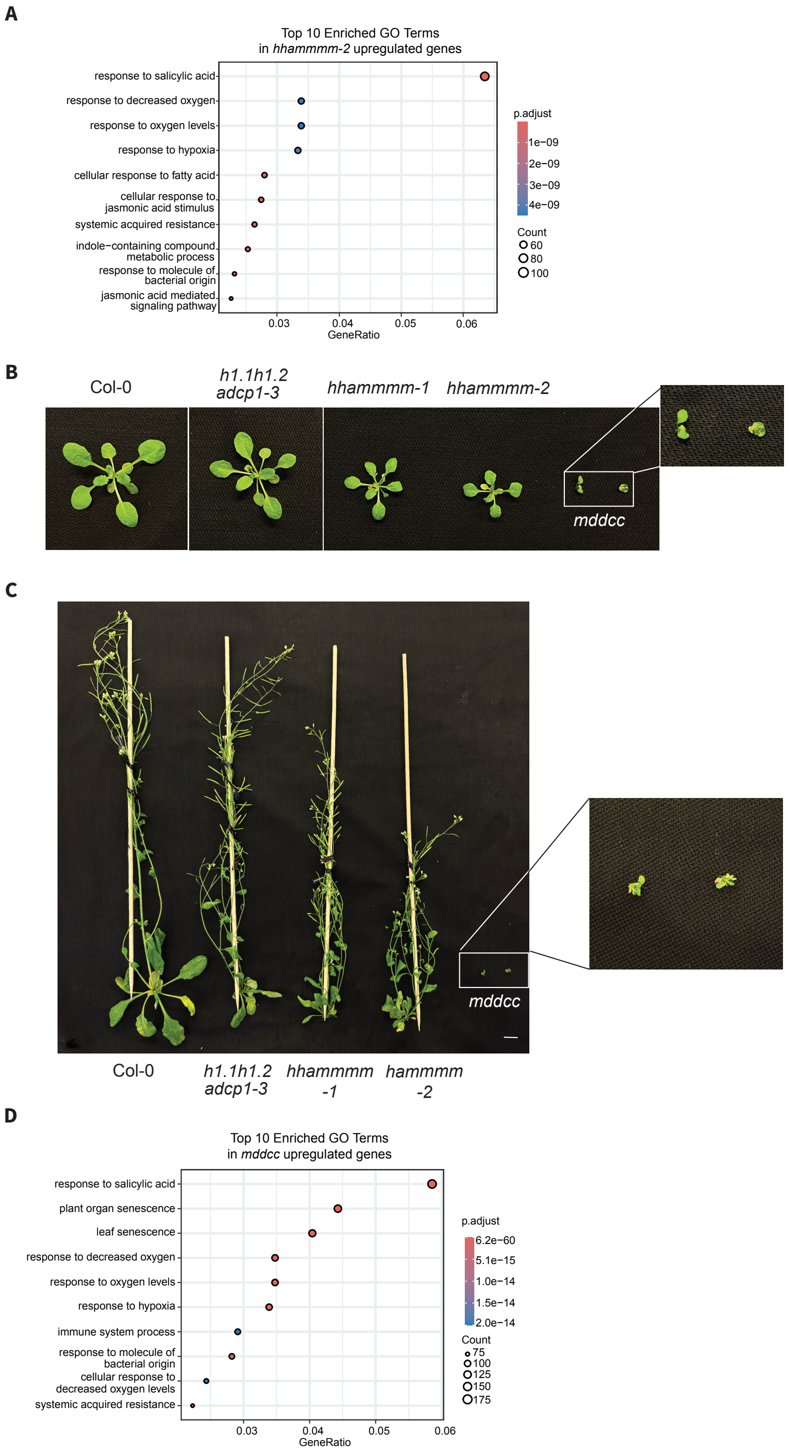


**Fig. S4. Phenotypes of the *hhammmm* mutants resemble classical autoimmune mutants.** **A.** Top 10 enriched GO terms of *hhammmm-2* upregulated genes. **B.** Photos showing the morphology of 4-week-old seedlings of Col-0, *h1.1 h1.2 adcp1-3*, *hhammmm*, and *mddcc* mutants. **C.** Photos demonstrating the morphology of 6-week-old adult plants of Col-0, *h1.1 h1.2 adcp1-3*, *hhammmm*, and *mddcc* mutants. **D.** Top 10 enriched GO terms of *mddcc* upregulated genes.

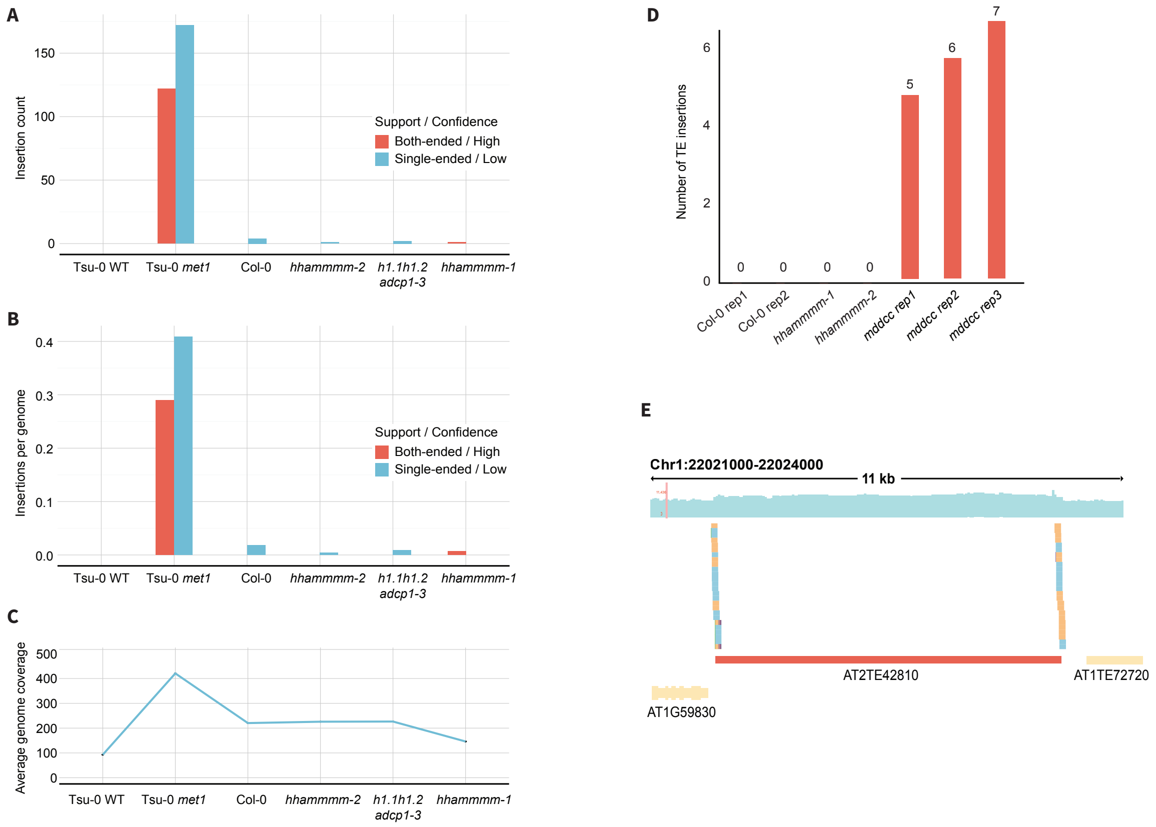


**Fig. S5. TE activation in the *hhammmm* mutants is insufficient for de novo transposition.** **A.** TE insertion count, **B.** TE insertion normalized by genome coverage, or **C.** average genome coverage among Tsu-0 WT, Tsu-0 *met1*, *h1.1 h1.2 adcp1-3*, and *hhammmm* mutants from Pacbio HiFi sequencing. **D.** Bar plot showing the number of TE insertions detected in Col-0, *hhammmm*, and *mddcc* mutants. **E.** IGV screenshot showing read alignments at the junction between the original genomic sequence and the inserted TE in the *mddcc* mutant. The green bar indicates the position of the TE insertion site.

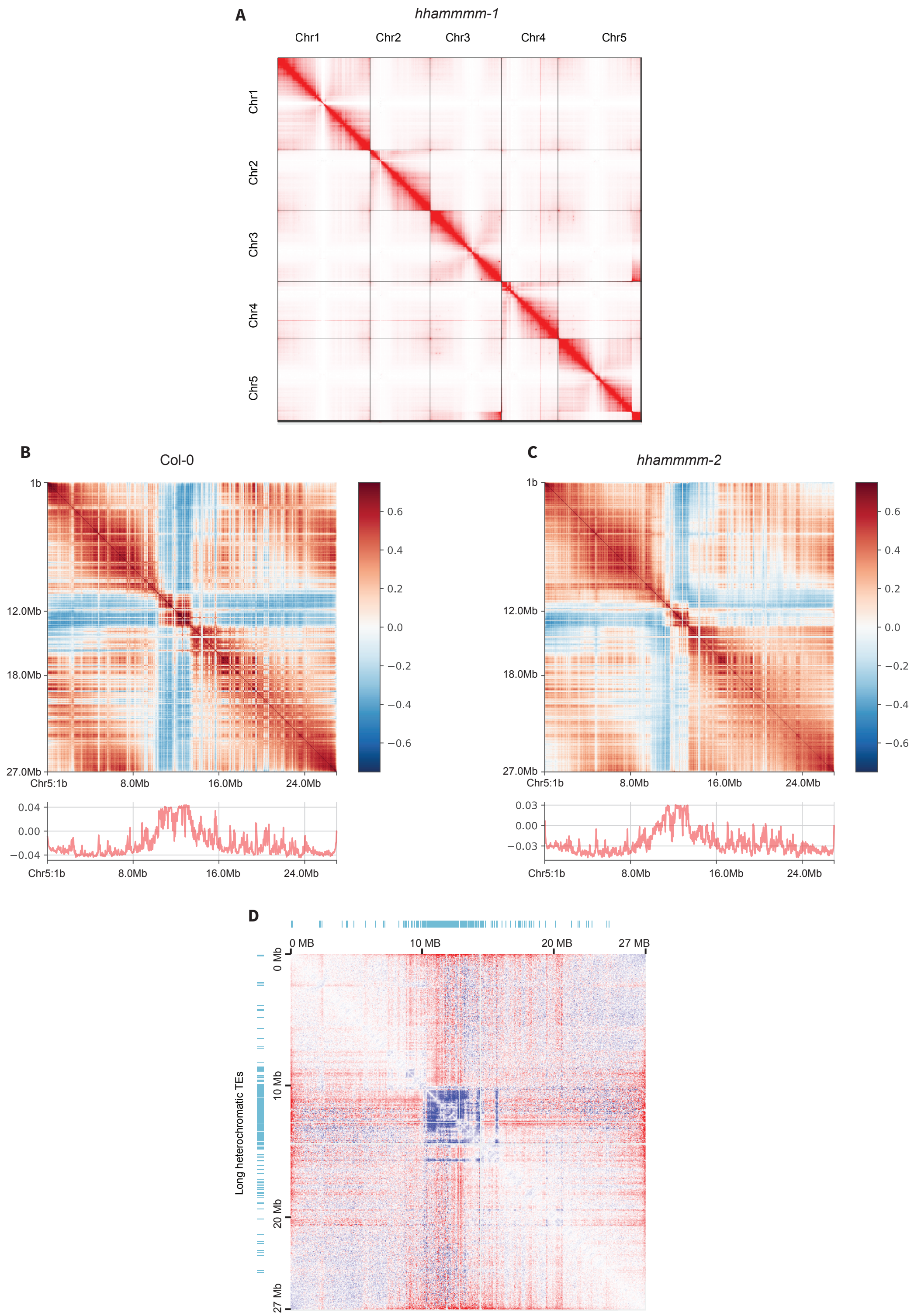


**Fig. S6. Chromosome rearrangement is observed in the *hhammmm-1* mutant.**
**A.** Genome-wide observed contact matrix of *hhammmm-1* mutant. The end of Chr5 is translocated to the end of Chr3. Square matrices with eigenvector plots showing the observed/expected interactions at Chr5 for **B.** Col-0, and **C.** *hhammmm-2* mutant. **D.** Micro-C map of Chr5 showing the differences in interactions between *hhammmm-2* and Col-0.

**Table S1. Micro-C replicates show data consistency and reproducibility.**
The table summarizes the data quality metrics of Col-0, *hhammmm-1,* and *hhammmm-2* replicates.

| Metric | Col-rep1 Value | Col-rep1 Percentage | Col-rep2 Value | Col-rep2 Percentage |
|---|---|---|---|---|
| Total Read Pairs | 342,615,927 | 100% | 410,532,607 | 100% |
| Unmapped Read Pairs | 64,705,558 | 18.89% | 64,576,573 | 15.73% |
| Mapped Read Pairs | 229,638,728 | 67.03% | 290,040,546 | 70.65% |
| PCR Dup Read Pairs | 86,699,940 | 25.31% | 111,699,861 | 27.21% |
| No-Dup Read Pairs | 142,938,788 | 41.72% | 178,340,685 | 43.44% |
| No-Dup Cis Read Pairs | 119,148,434 | 83.36% | 148,615,686 | 83.33% |
| No-Dup Trans Read Pairs | 23,790,354 | 16.64% | 29,724,999 | 16.67% |
| No-Dup Valid Read Pairs (cis >= 1kb + trans) | 112,854,050 | 78.95% | 146,428,386 | 82.11% |
| No-Dup Cis Read Pairs < 1kb | 30,084,738 | 21.05% | 31,912,299 | 17.89% |
| No-Dup Cis Read Pairs >= 1kb | 89,063,696 | 62.31% | 116,703,387 | 65.44% |
| No-Dup Cis Read Pairs >= 10kb | 66,410,478 | 46.46% | 86,654,253 | 48.59% |

| Metric | *hhammmm*-1-rep1 Value | *hhammmm*-1-rep1 Percentage | *hhammmm*-2-rep1 Value | *hhammmm*-2-rep1 Percentage |
|---|---|---|---|---|
| Total Read Pairs | 393,574,614 | 100% | 393,925,825 | 100% |
| Unmapped Read Pairs | 61,141,991 | 15.54% | 52,449,261 | 13.31% |
| Mapped Read Pairs | 278,908,949 | 70.87% | 289,484,430 | 73.49% |
| PCR Dup Read Pairs | 105,753,235 | 26.87% | 108,245,385 | 27.48% |
| No-Dup Read Pairs | 173,155,714 | 44.00% | 181,239,045 | 46.01% |
| No-Dup Cis Read Pairs | 144,942,180 | 83.71% | 152,401,414 | 84.09% |
| No-Dup Trans Read Pairs | 28,213,534 | 16.29% | 28,837,631 | 15.91% |
| No-Dup Valid Read Pairs (cis >= 1kb + trans) | 135,920,074 | 78.50% | 143,607,550 | 79.24% |
| No-Dup Cis Read Pairs < 1kb | 37,235,640 | 21.50% | 37,631,495 | 20.76% |
| No-Dup Cis Read Pairs >= 1kb | 107,706,540 | 62.20% | 114,769,919 | 63.33% |
| No-Dup Cis Read Pairs >= 10kb | 81,530,743 | 47.09% | 86,579,862 | 47.77% |
